# Antimicrobial peptides can generate tolerance by lag and interfere with antimicrobial therapy

**DOI:** 10.1101/2022.06.03.494509

**Authors:** Daniel Sandín García, Javier Valle, Jordi Morata, David Andreu, Marc Torrent Burgas

**Author notes:** Corresponding Authors: David Andreu; Marc Torrent.

## Abstract

Antimicrobial peptides (AMPs) are widely distributed molecules secreted mostly by cells of the innate immune system to prevent bacterial proliferation at the site of infection. As with classic antibiotics, continued treatment with AMPs can create resistance in bacteria. However, whether AMPs can generate tolerance as an intermediate stage towards resistance is not known. Here we show that treatment of *Escherichia coli* with different AMPs induces tolerance by lag, particularly for those peptides that have internal targets. This tolerance can be detected as different morphological and physiological changes, which depend on the type of peptide molecule the bacterium has been exposed to. In addition, we show that AMP tolerance can also affect antibiotic treatment. Genomic sequencing of AMP-tolerant strains shows that different mutations alter membrane composition, DNA replication, and translation. Some of these mutations have also been observed in antibiotic-resistant strains, suggesting that AMP tolerance could be a relevant step in the development of antibiotic resistance. Monitoring AMP tolerance is relevant with a view to eventual therapeutic use of AMPs and because cross-tolerance might favor the emergence of resistance against conventional antibiotic treatments.

## INTRODUCTION

Tolerant bacteria can emerge when cell cultures are periodically treated with antibiotics. The tolerant phenotype does not provide resistance to antibiotics, i.e., the antibiotic minimum inhibitory concentration (MIC) remains unchanged^1^. However, it does allow the cells to survive longer due to mutations that lengthen the latency state^2,3^. This tolerant state precedes resistance since the acquired mutations allow the bacteria to survive longer, increasing the likelihood of developing resistance. In fact, the tolerant state has been proposed as an intermediate step required for the ultimate development of resistance^4^. However, tolerance by lag has not yet been described for all antibiotics, particularly not for AMPs.

In the current context of increasingly resistant strains and scarcity of new antibiotics, AMPs are one of the most promising strategies to treat infections by antibiotic-resistant bacteria^5^. AMPs are widely distributed in nature and have a very potent activity and a broad mechanism of action^6^. Although AMP action is mainly associated with cell membrane disruption^7,8^, AMPs can also exert their action by interacting with internal targets, at many levels in targeted bacteria^9^. Although long-term treatment with AMPs can also lead to resistant strains, as found for colistin^10^, no studies address the development of tolerance and how it may affect the treatment of infections.

Here, we present evidence that AMP treatment can induce tolerance by lag, particularly when there is an internal AMP target. These tolerant strains have phenotypes very similar to those already described by antibiotics, with colony morphologies of reduced size, an increase in the latency phase, and less pronounced lethality curves. Our results show that these tolerant strains can hinder the action of other antimicrobials, including conventional antibiotics. In this context, we propose that tolerance by lag should be considered when developing new AMPs, as well as the collateral effects in the treatment of infections with antibiotics.

## RESULTS

### Antimicrobial peptides generate tolerance by lag

Tolerance by lag (henceforth tolerance) can emerge when bacterial cultures are periodically treated with antibiotic concentrations equivalent to or higher than the MIC for a limited time. In these conditions, some cells can survive the treatment and adjust to the stress conditions, later becoming tolerant and increasing the population survival rate. To determine the existence of tolerance in the case of AMPs, we selected four peptides with different mechanisms of action (Table 1). On the one hand, polymyxin B (PolB) and LL-37 because they display a classic mechanism of action, involving partial insertion into the membrane and subsequent depolarization leading to bacterial cell death^11,12^. On the other hand, pleurocidin (Pleu) and dermaseptin (Derm) as peptides that enter the bacterium cytoplasm and interact with internal targets, affecting essential metabolic processes^13,14^. Here, we incubated the cells with peptides at the MIC and optimized the incubation time for each peptide to ensure a survival percentage below 0.1% (Figure 1a).

**Table 1.**
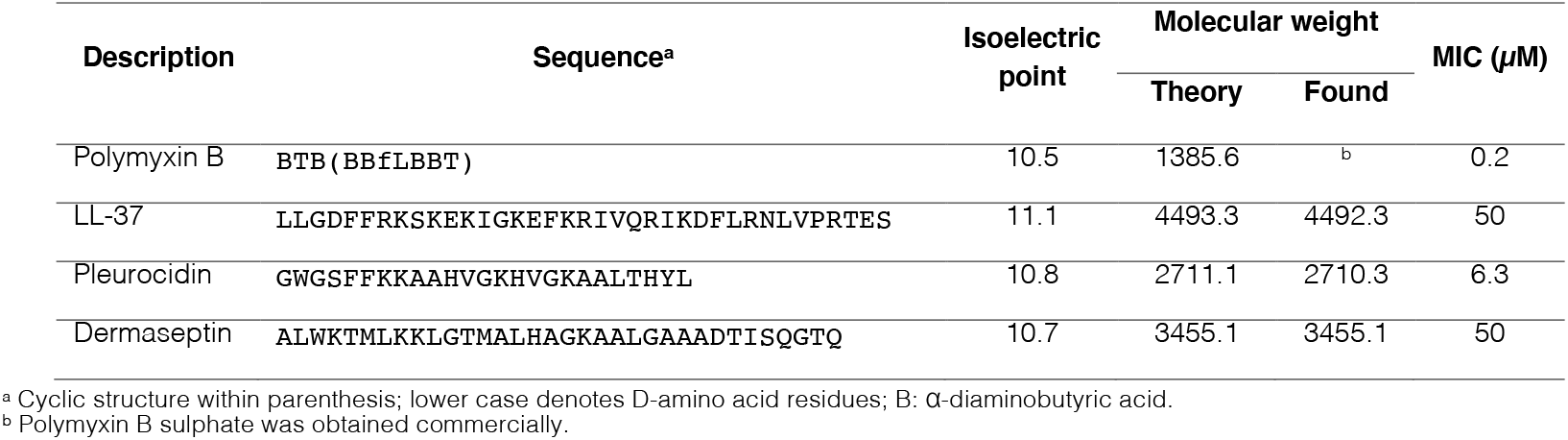
AMP sequences and MIC under assay conditions.

**Figure 1.**
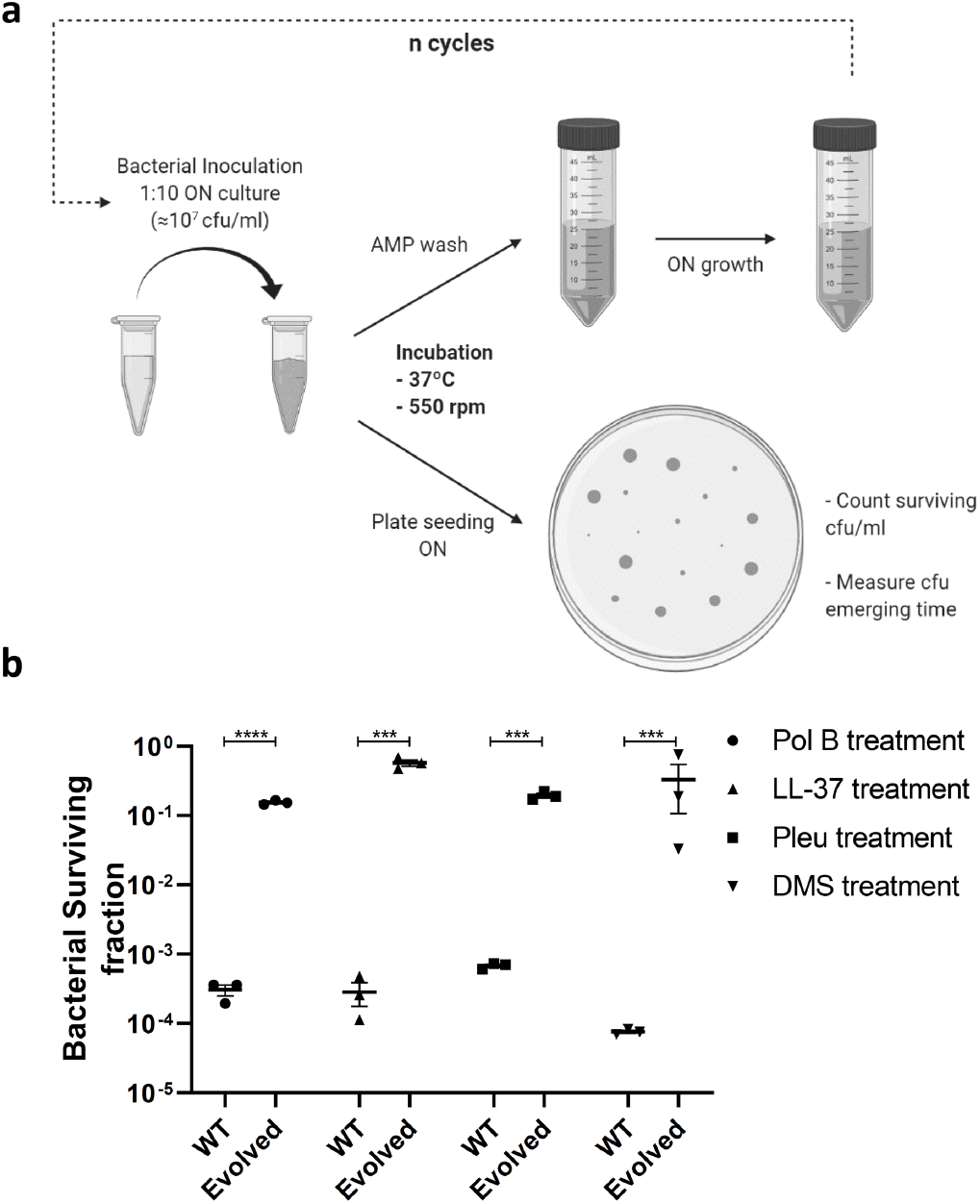
Periodic incubation of bacteria with AMPs. (a) Development of tolerant strains in *E. coli* against the tested AMPs. Cell cultures were evolved for 10 successive cycles or until resistance was detected (PolB, after 8 cycles) (b) Bacterial survival fraction before and after the assay. Values are shown as the mean ± SEM and individual replicates are displayed.

After 10 sequential cycles of evolution, an increase in cell survival was observed, with final values from 0.1% to 10% or even higher (Figure 1b). Under these conditions, resistance was only observed for PolB after 8 incubation cycles. This development of resistance was specific to this AMP and no cross-resistance was observed with other peptides (Supplementary Table 1).

After treating the bacterial cultures with AMPs, all cultures showed a characteristic decrease in colony size when surviving cells were plated in Petri dishes, except for LL-37 (Figure 2a,b). These results suggest that cells respond to the stress caused by AMPs and decrease their metabolism to become less susceptible, probably by increasing the lag time. However, this phenotype could be transient or could later be fixed in the genome to generate tolerance. One of the most distinctive features of tolerance is the increase in the lag phase after repeated cycles of incubation. To elucidate whether the treatment with AMPs produced similar characteristics, we measured the colony size distribution after each cycle of evolution using ScanLag. This method measures the distribution of lag times in individual cells from serial images of individual colonies in Petri dishes. In this way we were able to observe significant differences in colony size after AMP treatment (Figure 2c). Pleu displayed the largest effect, of comparable magnitude to ampicillin, as previously described^1^. These effects were observed even after the first incubation cycle, reaching maximum values after 3-4 cycles. Derm generated tolerance in later incubation cycles, and the effect, although milder compared to Pleu, was significant. On the other hand, the effect was almost zero for LL-37 and, for PolB, we even observed a decrease in the lag phase. While PolB could generate small colonies after the first incubation, the phenotype was not retained in later cycles, suggesting that the trait was not fixed in the genome. The observed results suggest that tolerance in AMPs appears to be indicative of intracellular targets, while membrane-level effects would not cause a measurable effect.

**Figure 2.**
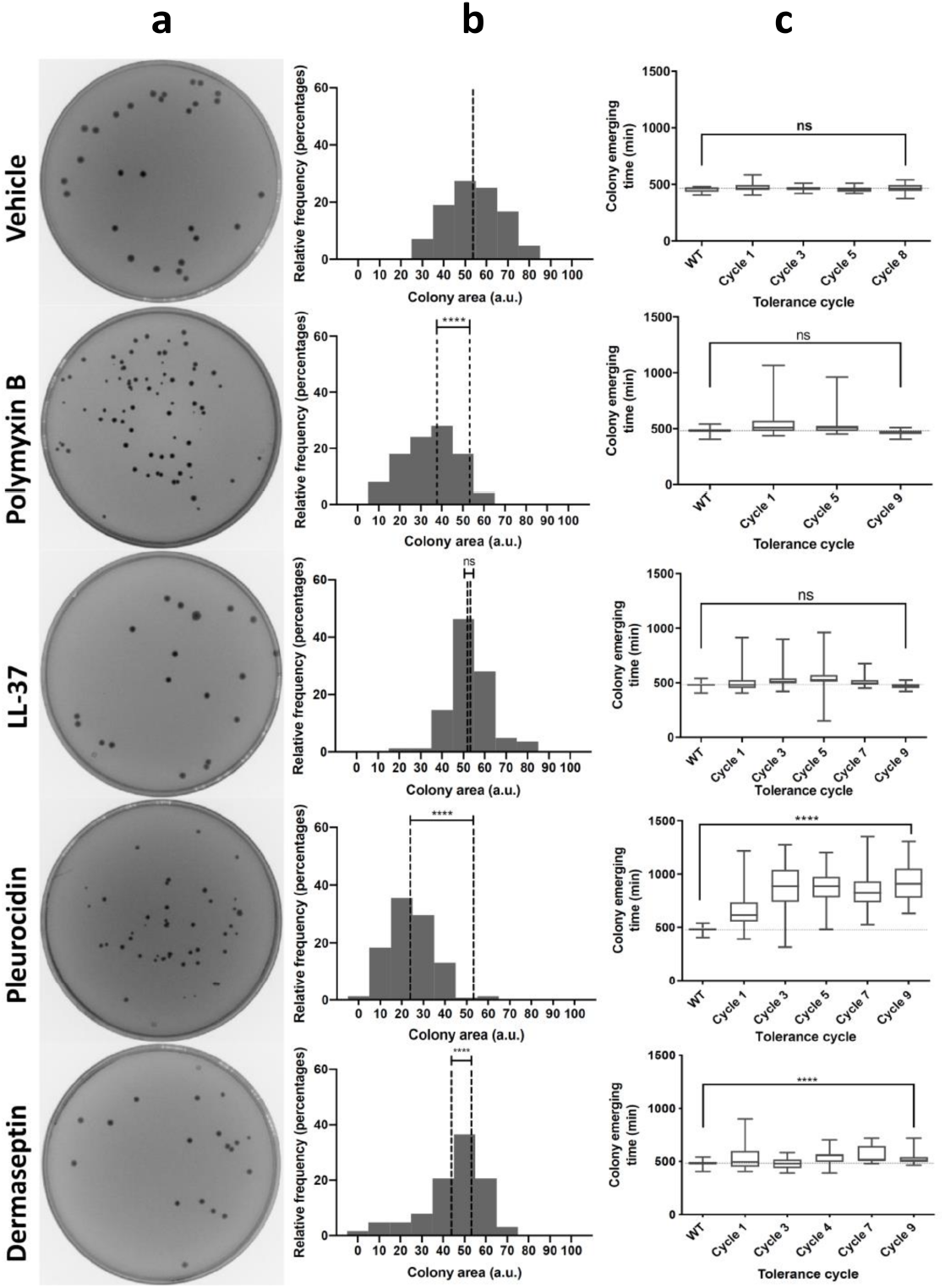
Development of tolerance after AMP treatment. (a) Plate images before and after the first treatment of bacteria with AMPs. (b) Colony area distribution (accumulated in 3 replicates) is shown beside the image plates. Treated and reference means are displayed as dotted lines. (c) Colony appearance times as measured with ScanLag vs. the different AMPs. The vehicle control is displayed for comparison. All experiments were done in triplicate.

### Tolerance in AMPs can affect antimicrobial treatments

After evolving *E. coli* cells tolerant to different AMPs, we investigated whether the strains were cross-tolerant to other AMPs or, on the contrary, the phenotype was AMP-specific. To this end, we determined the lethality curves for each of the evolved strains with all AMPs used in this study (Figure 3). We observed that, in general, strains evolved with an AMP developed tolerance against that same AMP. In line with ScanLag results, the Pleu-evolved strain was the one displaying a larger effect, which can be observed even after the second incubation cycle. For Derm, a significant tolerance was also observed, with a drop in the slope of the lethality curve from cycle 4 onwards. For LL-37 and PolB, we did not find significant changes in the lethality curves. In the case of PolB, it should be noted that resistance was detected in cycle 8, so the observed changes in the lethality curve should not be attributed to tolerance. In those cases where cross-tolerance to different AMPs was observed, the effects were generally small and, in most cases, not significant. The absence of cross-tolerance between Derm and Pleu suggests that, despite acting both on internal targets, the mechanism of action would be different. A slight cross-tolerance to LL-37 was observed in strains evolved with Derm, which could be due to non-specific changes at the cell membrane level.

**Figure 3.**
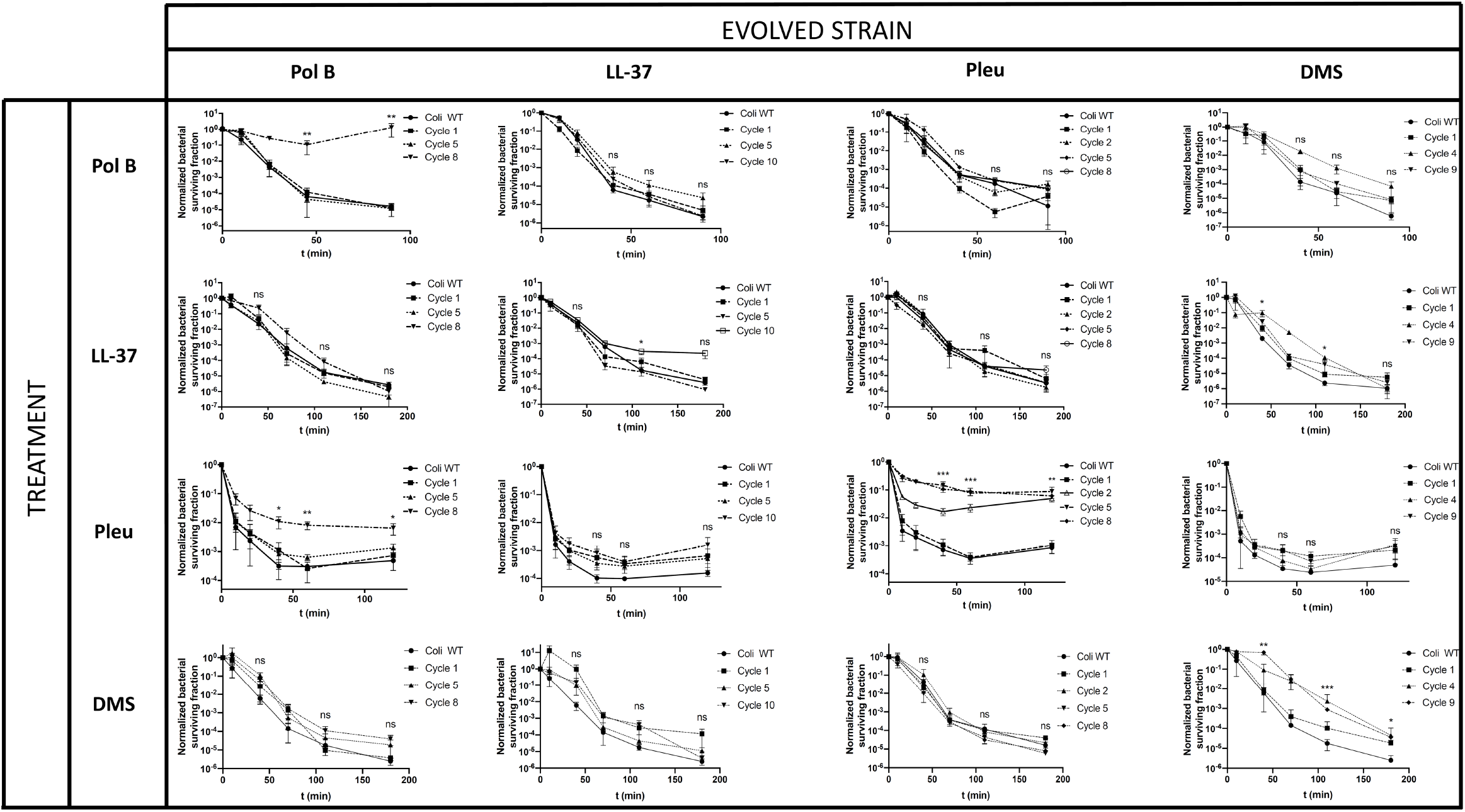
Killing curve assays for evolved strains against all AMPs tested. All combinations for evolved strains and AMPs were tested to detect cross-tolerance. The data is given as the relative surviving bacteria compared to the initial inoculum. Values are shown as the mean ± SEM, all experiments were performed in triplicates.

Finally, we investigated whether our evolved strains could display cross-tolerance with classical antibiotics. To this end, strains evolved with either LL-37 or Pleu were incubated with ampicillin (binds penicillin-binding proteins and inhibits cell wall synthesis), kanamycin (binds the ribosome, inhibiting translation), ciprofloxacin (binds topoisomerase IV, inhibiting replication), and nalidixic acid (binds DNA gyrase, inhibiting transcription and replication). In all cases, evolved strains displayed the same MIC against the antibiotics, compared with the parental strain, showing that they were not antibiotic-resistant (Supplementary Table 2). However, significant effects were observed in kanamycin action against strains evolved with Pleu, as detected by the killing curves (Figure 4). Interestingly, these effects sensitized *E. coli* to kanamycin, suggesting that, even if both antimicrobials can bind the ribosome, the site of action might be different. These findings are in tune with earlier observations that resistance to a given AMP could sensitize bacterial cells to other antimicrobials^15^. We also observed a significant difference in ciprofloxacin and nalidixic acid for Pleu-evolved strains, suggesting a pleiotropic binding to cytoplasmatic proteins (Figure 4). In these cases, the evolved strains displayed tolerance to antibiotics, reflected as slower killing curves. These results are also consistent with previous reports in which Pleu at high concentrations could inhibit transcription, translation, and replication in bacterial cells^16^.

**Figure 4.**
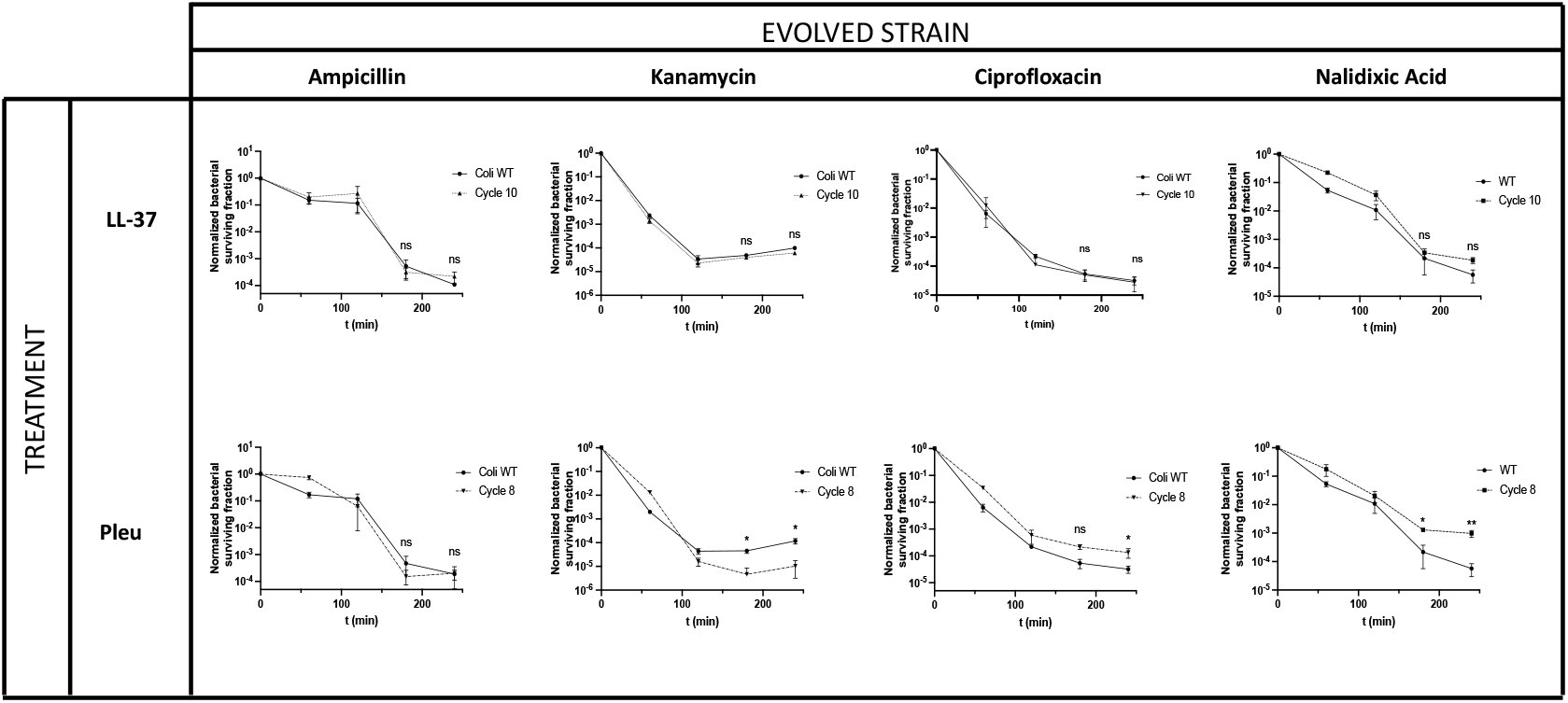
Killing curve assays after antibiotic exposure in evolved *E. coli* strains with Pleu and LL-37. Bacterial viability was measured as the relative number of surviving CFUs compared with the initial inoculum. The concentrations used for each antibiotic were 6.3 μg/mL for ampicillin, 12.5 μg/ml for kanamycin, 0.1 μg/mL for ciprofloxacin, and 20 μg/mL for nalidixic acid. Values are shown as the mean μ SEM, all experiments were performed in triplicates.

### The tolerance mutational landscape is diverse

We showed that AMP tolerance can be fixed in the genome, as we detected a significant change in the lethality curve for the evolved strains. To determine the mutations responsible for that tolerance, we sequenced (by duplicate) the genome of both Pleu- and LL-37-evolved strains. For Pleu, we detected in all reads and both replicates, only one conserved change corresponding to the deletion of an adenosine (A) nucleotide in an intergenic region containing the promoter for the yejK and yejL genes (Table 2). The first gene encodes for radD helicase while the second is uncharacterized. These mutations could explain the increased tolerance to antibiotics of the quinolone family, such as ciprofloxacin and nalidixic acid, involved in the replication of the bacterial genome. We also found other mutations (≥90% in the population) involving several genes related to the ribosome (sspA), glycolate metabolism (glcE), and phospholipid transport in the membrane (mlaC-mlaF, gltP; Table 2). The first two types (related to ribosome and glycolate metabolism) could explain the sensitization effect observed against kanamycin. In the first case, because the ribosome is the main target of kanamycin, and in the second because it has been shown that metabolism linked to carbon sources allows bacterial sensitization against amino acids of the aminoglycoside family such as kanamycin^17^. For their part, changes at the membrane level, particularly in glycolipid transport (gltP), may result in asymmetry of lipid composition^18^ and thus explain the cross-tolerance observed between Pleu and LL-37.

**Table 2.**
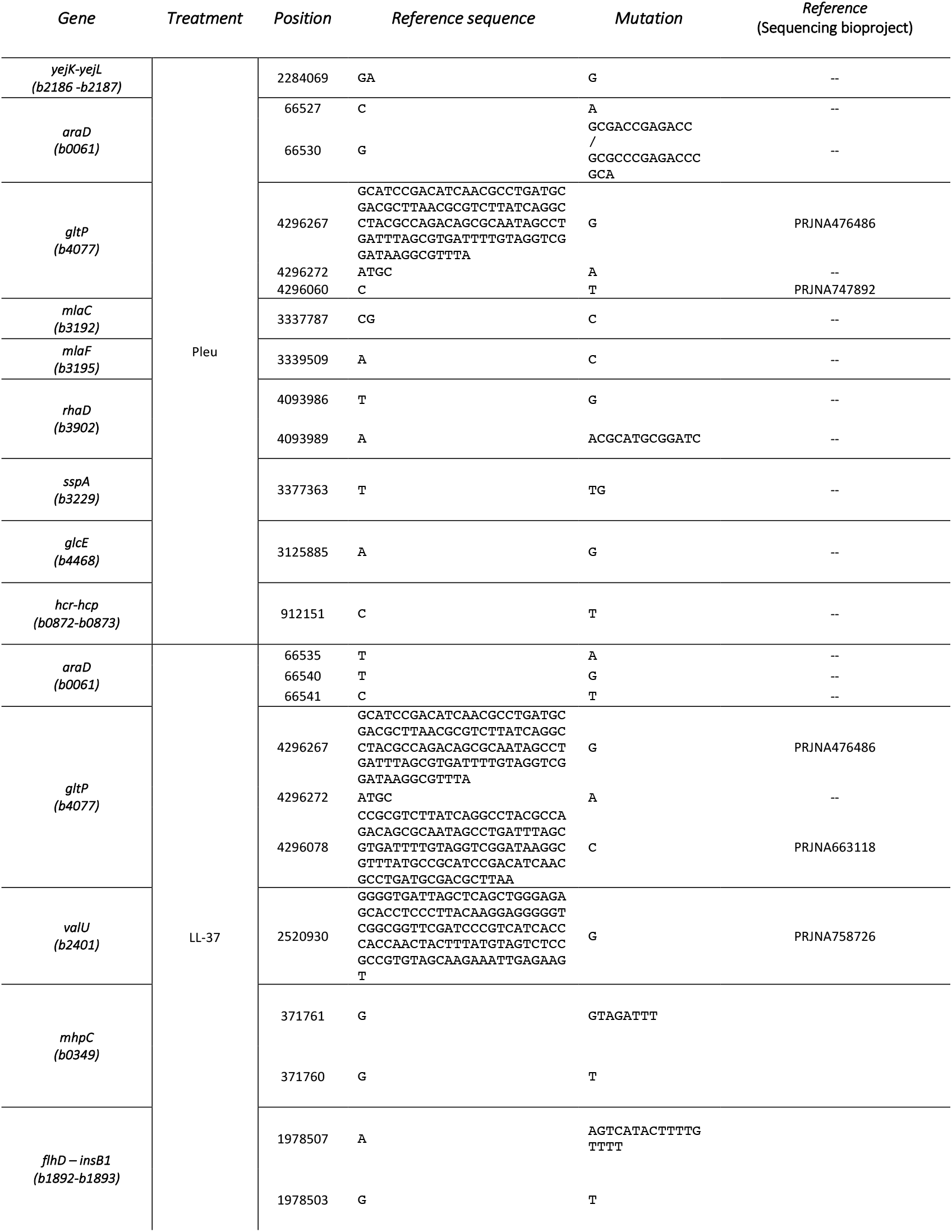
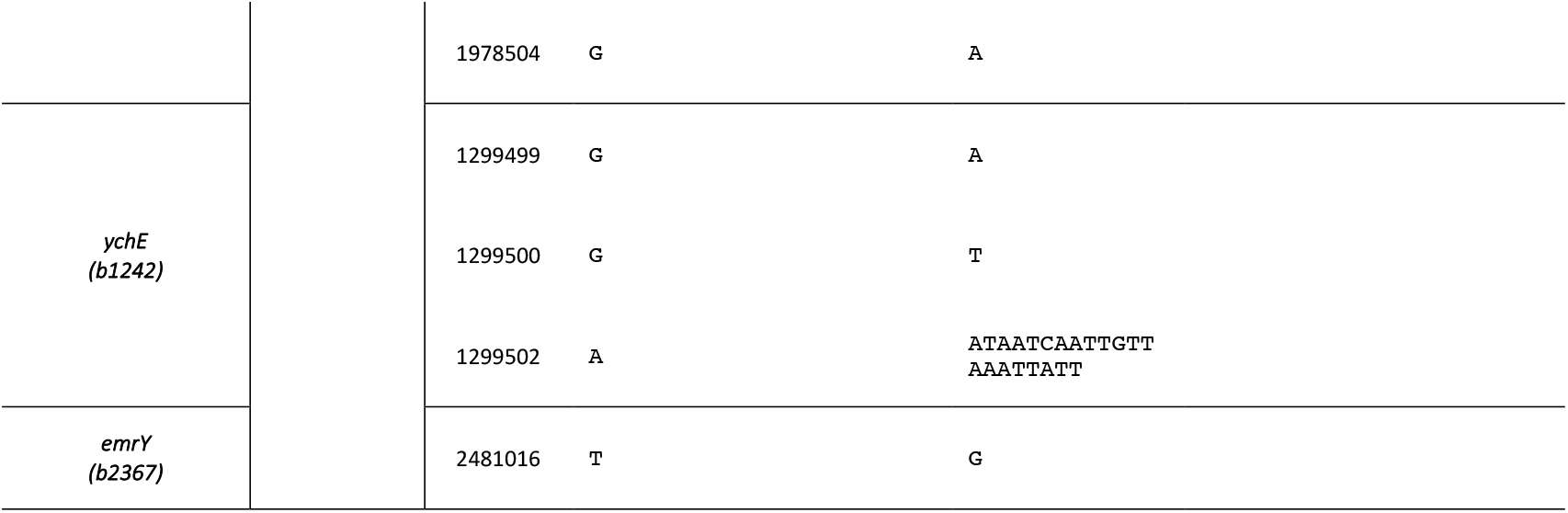
Summary of relevant mutations in Pleu and LL-37 evolved strains.

In the LL-37-evolved strain, we also found changes at the genomic level, although not directly related to internal, essential targets. Instead, they were associated to amino acid metabolism (mhpC, valU) and glycolipid transport (gltP), in line with the LL-37 mechanism of action. The absence of significant mutations in the evolved strains may explain the lack of tolerance after LL-37 treatment.

## DISCUSSION

The failure of antibiotic treatments is mainly attributed to the resistance acquired by microorganisms^19^. However, there are other relevant mechanisms including tolerance, that can contribute to therapeutic failure^20^. Antimicrobial tolerance refers to the ability of bacteria to survive the effect of the antimicrobial for longer periods. In antibiotics, tolerance can precede resistance as it allows the cell to survive for the time needed to develop resistance mutations^4^. These mechanisms acquire a special relevance in the case of AMPs, which are produced by the host innate immune cells and represent the first barrier of protection against infections^21^. Neutrophil degranulation, for example, involves the secretion of AMPs that reach high local concentrations in the body^22^. Therefore, ideal circumstances concur for bacterial cells to acquire mutations conferring some tolerance to these AMPs. In this study we have shown that bacteria can gain tolerance in the presence of high AMP concentrations (∼MIC)

This behavior is not identical for all AMPs. Some generated tolerance in early incubation cycles (1-4) while others failed to induce tolerance. While AMPs share some features, they are quite different in terms of sequence and structure. In the present case, all peptides are cationic, with an isoelectric point between 10 and 11, and adopt helical structure except PolB which is cyclic. Hence, the gain of tolerance cannot be directly related to structure. Based on our results, we hypothesize that the mechanism of action could explain the observed differences. Thus, both LL-37 and PolB act preferentially on the cell membrane while Pleu and Derm can inhibit the synthesis of macromolecules, including DNA, RNA, and proteins. The fact that Pleu-tolerant strains have altered sensitivity to kanamycin, ciprofloxacin, and nalidixic acid but not ampicillin, reinforces the idea of action at the level of cytoplasmic targets. Indeed, the action of both Pleu and Derm is more similar to classic antibiotics such as kanamycin, which would explain their behavior.

The increase in lag phase for tolerant strains suggests that bacteria can achieve tolerance after repeated AMP incubation. The appearance at the first incubation cycle of smaller colonies that are not actually tolerant suggests a two-stage mechanism. First, stress causes a decrease in the metabolic activity of bacteria, which translates into smaller colonies and slower growth. Then, after several cycles of incubation, these changes can be fixed in the genome, generating tolerant strains. In the first few cycles, we observed colonies with reduced growth, probably because of transient heritable phenotypes at the proteome or epigenome level. Then, mutations at the genome level could accumulate in subsequent cycles, generating tolerant strains. This is well illustrated in the case of Derm, where colonies with reduced morphology were observed after the first incubation cycle but did not translate into tolerance as revealed by the lethality curves. However, from cycle 4 onwards, the lethality curves reflected the appearance of tolerance, suggesting mutations at the genome level.

The mutations observed in LL-37- and Pleu-evolved strains are consistent with the results observed in previous experiments. Some of these mutations were also observed in clinical strains, suggesting that tolerance can also happen in human patients (Table 2). It is tempting to speculate that, after the initial innate immune response, bacteria can become tolerant to AMPs. Then, when patients are treated with antibiotics, AMP tolerant strains may have an advantage in surviving antibiotics. As mutations allow them to survive longer times in the presence of antibiotics, resistance would be more likely to occur. The opposite behavior, sensitization to antibiotics, cannot be ruled out in view of our results, as noted by strains tolerant to Pleu, which were more sensible to kanamycin.

In summary, we show here that bacteria can generate tolerance against certain AMPs, depending on their mechanism of action. This tolerance can affect the action of other AMPs or even antibiotics used to treat these infections. Hence, the relationship between AMP activity and tolerance deserves further attention. On the one hand, a more comprehensive view of bacterial evolution to AMPs, involving both tolerance and resistance, could be useful when developing novel AMPs. On the other hand, further exploration of the mechanisms of cross-tolerance between antibiotics and AMPs should contribute to a better understanding of how bacteria become resistant to antibiotic treatment.

### EXPERIMENTAL SECTION

#### Materials

*Escherichia coli* strain was purchased from the Coli Genetic Stock Center (BW25113). All peptides but PolB (Thermo Fisher, Hampshire, UK) were synthesized by Fmoc solid-phase synthesis, as previously described^23^. Ampicillin was purchased from Merck (Darmstadt, Germany), kanamycin from Apollo Scientific (Stockport, UK), ciprofloxacin and nalidixic acid from Thermo Fisher (Hampshire, UK).

#### Peptide tolerant strain development and data processing

Overnight *E. coli* cultures were grown in LB medium and inoculums were prepared by dilution at 1:100 in fresh LB media (500 μL total volume). The cultures were incubated at 37ºC and 600 rpm in an Accutherm Microtube Shaking Incubator (Labnet International; Edison, NJ) at the corresponding MIC concentration (Table 1) for the corresponding time to ensure < 0.01% survival. Exposure times were 40, 30, 60, and 80 min for PolB, Pleu, LL-37, and Derm, respectively. After incubation, a sample of 400 μL from each assay was resuspended in 8 mL of fresh LB media to regrow the bacteria for the next cycle. The remaining volume (100 μL) was plated on Petri dishes. Plates were scanned every 15 min in an Epson Perfection V200 Photo instrument (Epson, Nagano, Japan) to monitor colony appearance time. Lag times were calculated from serial images using the ScanLag script, as previously described^24^. To determine colony size after the first incubation cycle, surviving cells after AMP treatment were plated on Petri dishes and allowed to grow overnight. Then, agar plates were scanned, and colony size was calculated using ImageJ.

#### Minimum inhibitory concentration (MIC)

Detection of bacterial growth inhibition was performed as previously described^25^, using a standard dilution at 5·10^5^ CFU/mL in Muller-Hinton medium. The MIC was defined as the last antimicrobial concentration where no visual growth could be detected.

#### Killing curve assay

Bacterial killing curves were measured by colony count in Petri plates at defined time points. Samples (500 μL, containing 450 μL of 1:100 bacteria overnight culture in LB media and 50 μL of 10x MIC AMP stocks) were taken at different times during the incubation with peptides and plated to Petri plates. After overnight incubation at 37ºC, CFUs were counted and survival rates at each point were calculated in comparison with an initial inoculum.

#### DNA Purification and genome sequencing

QIAamp® DNA Mini Kit (Qiagen, Germantown, MD) was used to isolate the *E. coli* whole genome in the wild-type strain and evolved strains following the manufacturer’s instructions. Purified whole genomes were sequenced with Oxford Nanopore (Oxford, UK).

Reads were mapped to *E. coli* BW25113 ASM75055v1 with BWA-MEM 0.7.17^26^. Alignment files containing only properly paired, uniquely mapping reads without duplicates were processed using Picard (http://broadinstitute.github.io/picard/) to add read groups and to remove duplicates. The Genome Analysis Tool Kit (GATK 4.1.8.0) HaplotypeCaller^27^ was used for variant calling. Joint genotyping was performed with combined gvcfs. Best Practices GATK filters were applied and variants with at least one sample containing the variant with read depth of at least 5 were retained. CNV prediction was performed with ControlFREEC 11.5^28^, using a pool of samples as CNV baseline and using windows of 20kb and 50kb. CNV calls from all samples that were less than 10kb apart were merged with Survivor^29^.

## ASSOCIATED CONTENT

Supporting information

## AUTHOR INFORMATION

## AUTHOR CONTRIBUTIONS

DA and MT designed, directed, obtained funding for, and coordinated the study. JV performed the synthesis of peptides and DS performed the experiments and JM performed the genome sequencing analysis. DS wrote an initial version of the paper, subsequently edited by DA and MT. All authors contributed to editing and revising the final version of the manuscript.

## ACKNOWLEDGMENT

The genome sequencing experiments were performed in the Centro Nacional de Análisis Genómico, a node of the Spanish Large-Scale National Facility ICTS.

## ABBREVIATIONS

AMP: antimicrobial peptide
PolB: polymyxin B
Pleu: pleurocidin
Derm: dermaseptin
MIC: minimum inhibitory concentration
CFU: colony-forming units

