## Supplementary_Information for "Antimicrobial peptides can generate tolerance by lag and interfere with antimicrobial therapy"

**Supplementary Table 1.** MICs in *E. coli* wild type and PolB resistant strain. MIC values are in μM.

| Peptide/Strain | *E. coli* wt | PolB |
|---|---|---|
| Pleurocidin | 0.06 | 0.06 |
| Polymyxin B | 0.01 | > 0.2 |
| Dermaseptin | 2 | 2 |
| LL-37 | 3 | 3 |

**Supplementary Table 2.** MICs in *E. coli* wild type and evolved. MIC values are in μM.

| | *E. coli* wt | PolB | Pleu | LL-37 | DMS |
|---|---|---|---|---|---|
| Ampicillin | 17 | 17 | 17 | 17 | 17 |
| Kanamycin | 1.4 | 1.4 | 1.4 | 1.4 | 2.8 |
| Ciprofloxacin | 42 | 42 | 42 | 42 | 42 |
| Nalidixic Acid | 11 | 11 | 11 | 11 | 11 |
